# Hypoxia-induced epigenetic silencing of polo-like kinase 2 promotes fibrosis in atrial fibrillation

**DOI:** 10.1101/445098

**Authors:** Stephan Reinhard Künzel, Karolina Sekeres, Susanne Kämmerer, Tomasz Kolanowski, Stefanie Meyer-Roxlau, Christopher Piorkowski, Sems Malte Tugtekin, Stefan Rose-John, Xiaoke Yin, Manuel Mayr, Jan Dominik Kuhlmann, Pauline Wimberger, Konrad Grützmann, Natalie Herzog, Jan-Heiner Küpper, Kaomei Guan, Michael Wagner, Ursula Ravens, Silvio Weber, Ali El-Armouche

**Affiliations:** Institute of Pharmacology and Toxicology, Faculty of Medicine Carl Gustav Carus, Technische Universität Dresden, Dresden, Germany; Department of Electrophysiology, Heart Center Dresden GmbH, Dresden, Germany; Department of Cardiac Surgery, Heart Center Dresden GmbH, Technische Universität Dresden, Dresden, Germany; Unit for Degradomics of the Protease Web, Institute of Biochemistry, University of Kiel, Kiel, Germany; The James Black Centre, King’s College, University of London, London, UK; Department of Gynecology and Obstetrics, Medical Faculty and University Hospital Carl Gustav Carus, Technische Universität Dresden, Germany; German Cancer Consortium (DKTK), Dresden and German Cancer Research Center (DKFZ), Heidelberg, Germany; National Center for Tumor Diseases (NCT), Partner Site Dresden, Germany; Brandenburg University of Technology, Senftenberg, Germany; Institut für Experimentelle Kardiovaskuläre Medizin, Universitäts Herzzentrum, Freiburg Bad Krotzingen, Freiburg im Breisgau, Germany

**Keywords:** Atrial fibrillation, fibroblasts, fibrosis, polo-like kinase 2, DNA methylation, osteopontin

## Abstract

Fibrosis and inflammation promote atrial fibrillation (AF) and worsen its clinical outcome. The underlying molecular mechanisms, that are relevant for effective antifibrotic drug development, are still under debate. This study deciphers a novel mechanistic interplay between polo-like kinase 2 (PLK2) and the pro-inflammatory cytokine osteopontin (OPN) in the pathogenesis of atrial fibrosis. Compared to sinus rhythm (SR) controls, right atrial appendages and isolated right atrial fibroblasts from AF patients showed downregulation of *PLK2* mRNA and protein levels, which were accompanied by remarkable hypoxia-sensitive DNA-methylation of the *PLK2* promotor. In an experimental setting, both, genetic deletion and pharmacological inhibition of PLK2 induced myofibroblast differentiation and reduced fibroblast proliferation. Notably, proteomics from *PLK2*-deleted fibroblasts revealed *de novo* secretion of OPN. Accordingly, we observed higher OPN plasma levels in AF patients with atrial fibrosis compared to non-fibrosis AF patients. Hence, we provide evidence for PLK2 reactivation and/or OPN inhibition as potential novel targets to prevent fibrosis progression in AF.

## Introduction

Atrial fibrillation (AF) is the most prevalent arrhythmia in the clinical routine. Reoccurring AF episodes lead to marked changes in the cardiac tissue architecture (Harada *et al*, 2015; Hadi *et al*, 2010; Boos *et al*, 2006) and contribute to morbidity and mortality. Current pharmacological therapies aim predominantly at directly suppressing arrhythmia by ion channel blockade. However, these approaches tend to be insufficient and tend to produce detrimental side effects that include arrhythmias (Heijman *et al*, 2018). Recently, there is paradigm change in AF pathophysiology pointing to AF being a (systemic) inflammatory disease that is not confined to the atria. Therefore, we chose to examine a specific cell population, the cardiac fibroblasts, that are involved in inflammation and fibrosis. The approach to target specific inflammatory mediators and fibrosis-associated genes in fibroblasts may constitute a novel approach to prevent or treat the progression of AF (Rudolph *et al*, 2010).

Any kind of cardiac injury or disorder can activate fibroblasts leading to myofibroblast differentiation (Tallquist & Molkentin, 2017; Nattel *et al*, 2008). Once activated, myofibroblasts gain size, express orderly arranged filaments of alpha smooth muscle actin (αSMA) and become highly secretory, resulting in deposition of interstitial collagen as well as local enrichment of cytokines and other inflammation mediators (Tallquist & Molkentin, 2017; Baum & Duffy, 2011). There is strong phenomenological evidence for altered fibroblast function in persistent AF in terms of increased myofibroblast differentiation and reduced fibroblast proliferation (Poulet *et al*, 2016). The underlying molecular mechanisms still remain widely elusive.

A number of inflammation mediators like interleukins 2, 6, 8, and 10, C-reactive protein, TNF-α or matrix metalloproteinases 1 and 2 as well as osteopontin (OPN) have been associated with the pathogenesis and progression of AF. They are considered to induce the fibrotic and proarrhythmic substrate in AF patients (Calvo *et al*, 2018; Chung *et al*, 2001; Watanabe *et al*, 2005; Vílchez *et al*, 2014). Several of those biomarkers have been found in both cardiac tissue biopsies and the peripheral blood of AF patients (Harada *et al*, 2015; Vílchez *et al*, 2014).

Recent studies indicated promising potential of OPN as a therapeutic target in inflammation-associated cardiac disease (Zhao *et al*, 2016). OPN is a glycoprotein of the extracellular matrix secreted by macrophages, T-cells and fibroblasts in the heart. Latest studies have demonstrated cytokine-like functions of OPN, mediating inflammation, heart remodeling processes, fibrosis and increasing the risk of atherosclerosis and heart failure (Zhao *et al*, 2016; İçer & Gezmen-Karadağ, 2018; López *et al*, 2013; Rubiś *et al*, 2018).

Recent research in our laboratory focused on differentially regulated genes in fibroblasts isolated from patients with persistent AF. An Affymetrix^®^ microarray (Poulet *et al*, 2016) revealed negative regulation of polo-like kinase 2 (PLK2) transcripts. PLK2 is a serine-threonine kinase that regulates cell cycle progression by centriole duplication and it is strongly associated with cell proliferation, mitochondrial respiration and apoptosis (Mochizuki *et al*, 2017; Li *et al*, 2014; Ma *et al*, 2003, 2). Published data suggest that PLK2 suppression may induce cellular senescence (Deng *et al*, 2017), a condition in which cells stop to proliferate (Coppé *et al*, 2008).

Thus, the objective of the current study was to clarify the (patho)physiological role of PLK2 in fibrosis, inflammation and the pathogenesis of permanent AF. we provide first evidence for a molecular link of dysregulated PLK2 expression and altered OPN secretion in pathological fibrosis.

## Results

### PLK2 expression is reduced in AF

Based on lower *PLK2* mRNA expression in AF fibroblasts in the Affymetrix^®^ microarray (Poulet *et al*, 2016), we confirmed a ~50% downregulation of *PLK2* mRNA abundance in fibroblasts isolated from AF patients as determined by quantitative real-time PCR (Figure 1 a). Furthermore, PLK2 protein expression was significantly reduced in human right atrial tissue samples from AF patients in comparison to those from patients with sinus rhythm (SR) (Figure 1 b and c) as determined by western blotting. Previous studies in non-cardiac tissues (Syed *et al*, 2006; Benetatos *et al*, 2011) have shown that PLK2 expression is strongly regulated by promoter hypermethylation. Therefore, we performed methylation-specific PCR for the *PLK2* promoter, which was positive in 6 out of 13 AF samples, while there was no detectable methylation of the *PLK2* promoter in any of the SR samples (n = 11) tested in our study (Figure 2 a and b). This suggests that AF pathogenies and/ or progression may involve epigenetic modifications affecting the *PLK2* promoter. To test this hypothesis, we exposed fibroblasts to chronic hypoxia (1% O_2_) treatment for 24, 72 and 96 h (Figure 2 c and d) since tissue hypoxia is present in AF and known to induce several epigenetic alterations (Gramley *et al*, 2010; Watson *et al*, 2014). After 24 h we found a 3.2-fold downregulation of *PLK2* mRNA expression (Figure 2 c). Methylation of the *PLK2* promoter was detectable after 96 h of chronic hypoxia treatment. As an additional positive control, cells were treated with 0.25 mM Dimethyloxaloylglycine (DMOG) which is an inhibitor of PHD finger protein (PHF) and factor inhibiting HIF (FIH-1) mimicking hypoxia by upregulation of hypoxia-inducible factor (HIF-1α) (Ayrapetov *et al*, 2011). 96 h DMOG treatment led to clearly detectable PLK2 promoter methylation.

**Figure 1:**
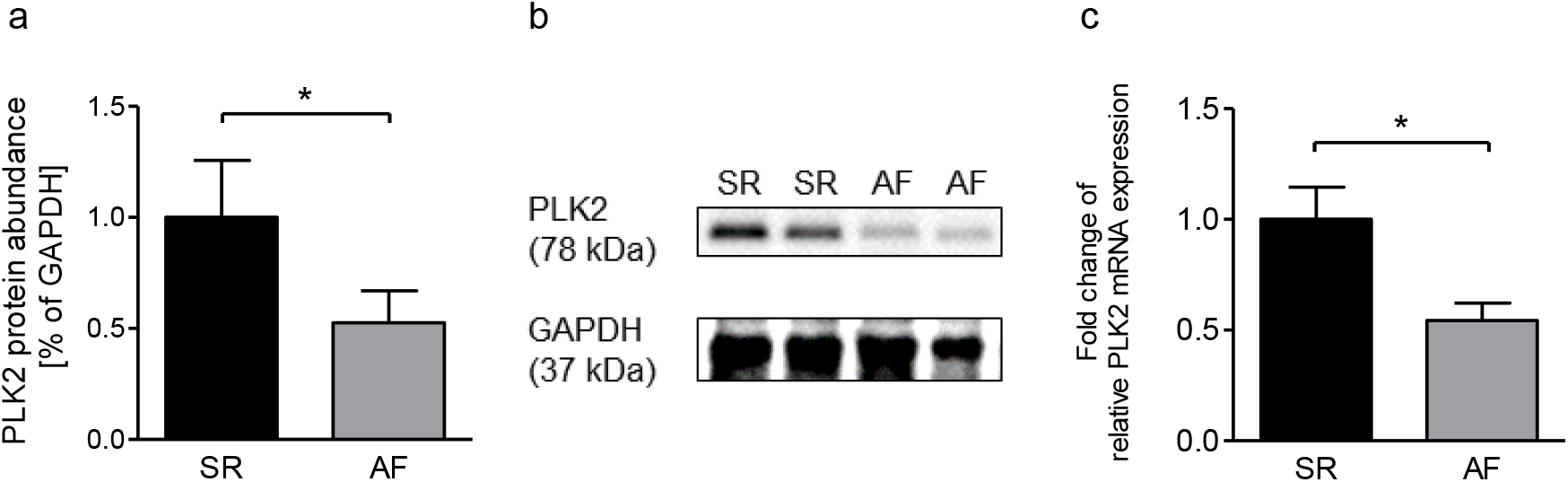
PLK2 is downregulated in patients suffering from atrial fibrillation. **a)** Expression of *PLK2* mRNA normalized to *GAPDH* in primary human atrial fibroblasts from SR and AF patients, analyzed with qPCR (n = 7 (SR) vs. 5 (AF)). **b)** Quantification of PLK2 protein abundance in human right atrial tissue samples from SR and AF patients analyzed by western blot (n = 9 per group). **c)** Representative western blot for b. ^*^: p < 0.05. SR: Sinus rhythm patients, AF: Atrial fibrillation patients.

**Figure 2:**
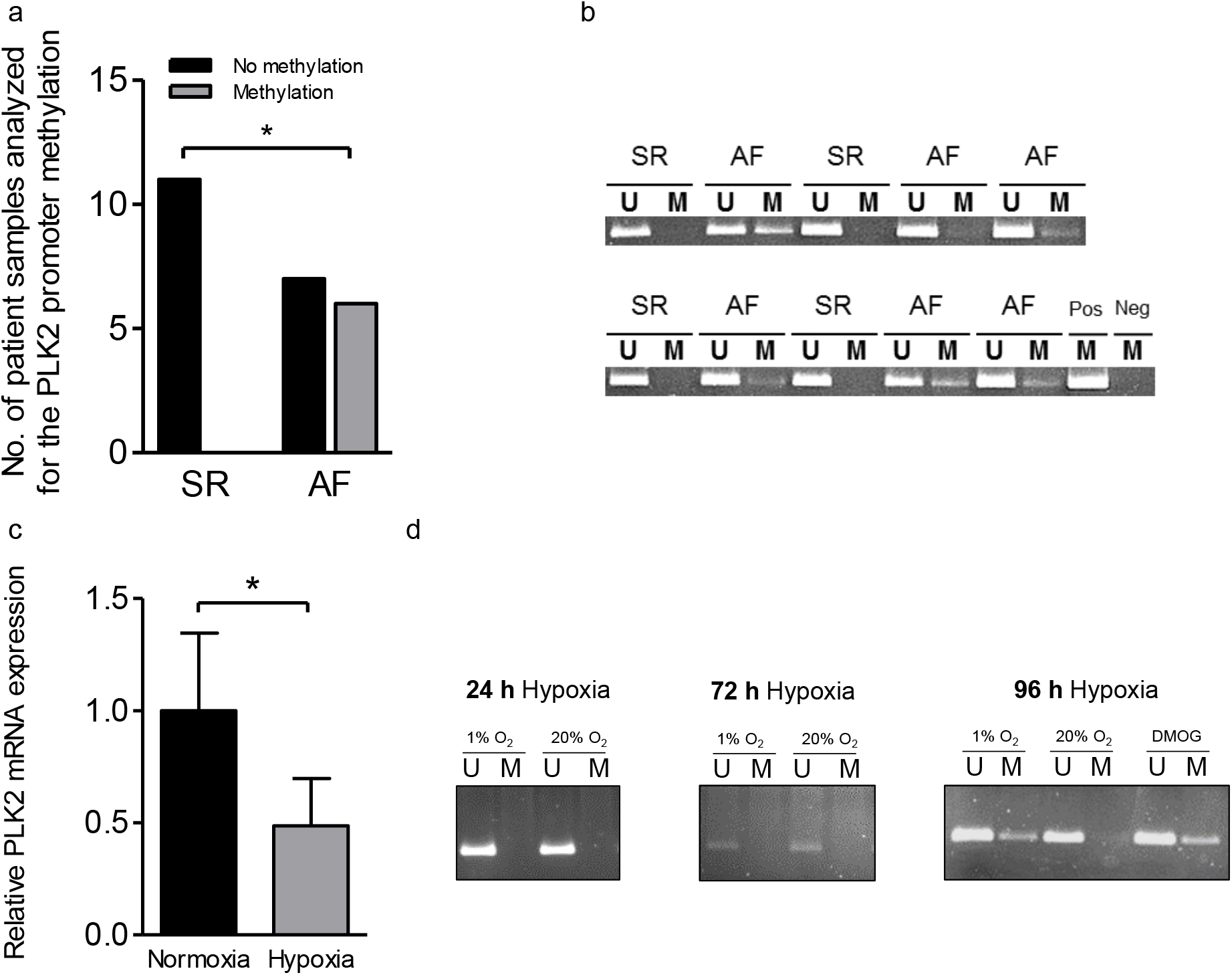
The *PLK2* promoter methylation status and *PLK2* expression are sensitive to hypoxia. **a)** Quantification of SR and AF heart tissue samples in which methylation was present or absent. Statistical analysis was done with Fisher‘s exact test (n_SR_ = 11; n_AF_ = 13). **b)** Gel images of a methylation-specific PCR of the *PLK2* promotor region (U: unmethylated, M: methylated, Pos: positive control (human universal methylated DNA standard), Neg: water control). **c)** Hypoxia-dependent (1% O_2_) expression of *PLK2* mRNA normalized to *EEF2* in human ventricular fibroblasts, analyzed with qPCR (n = 10 (Normoxia) vs. 7 (Hypoxia)). **d)** Gel images of a methylation-specific PCR of the PLK2 promotor region after 24 h, 72 h and 96 h of hypoxia treatment. Dimethyloxaloylglycine (DMOG) for 96 h was used as a positive control. (U: unmethylated, M: methylated, Pos: positive control (human universal methylated DNA standard), Neg: water control). ^*^: p < 0.05

### Inhibition and loss of PLK2 induces myofibroblast differentiation and abolishes fibroblast proliferation

Next, we studied the effects of pharmacological PLK2 inhibition (TC-S 7005, 1 μM, 7±1 days) in primary human SR fibroblasts and the consequences of genetic PLK2 deletion (PLK2 KO (Inglis *et al*, 2009)). TC-S 7005 and PLK2 KO markedly induced myofibroblast differentiation (Figure 3 a – c, and e – g) and reduced fibroblast proliferation rates (Figure 3 d and h).

**Figure 3:**
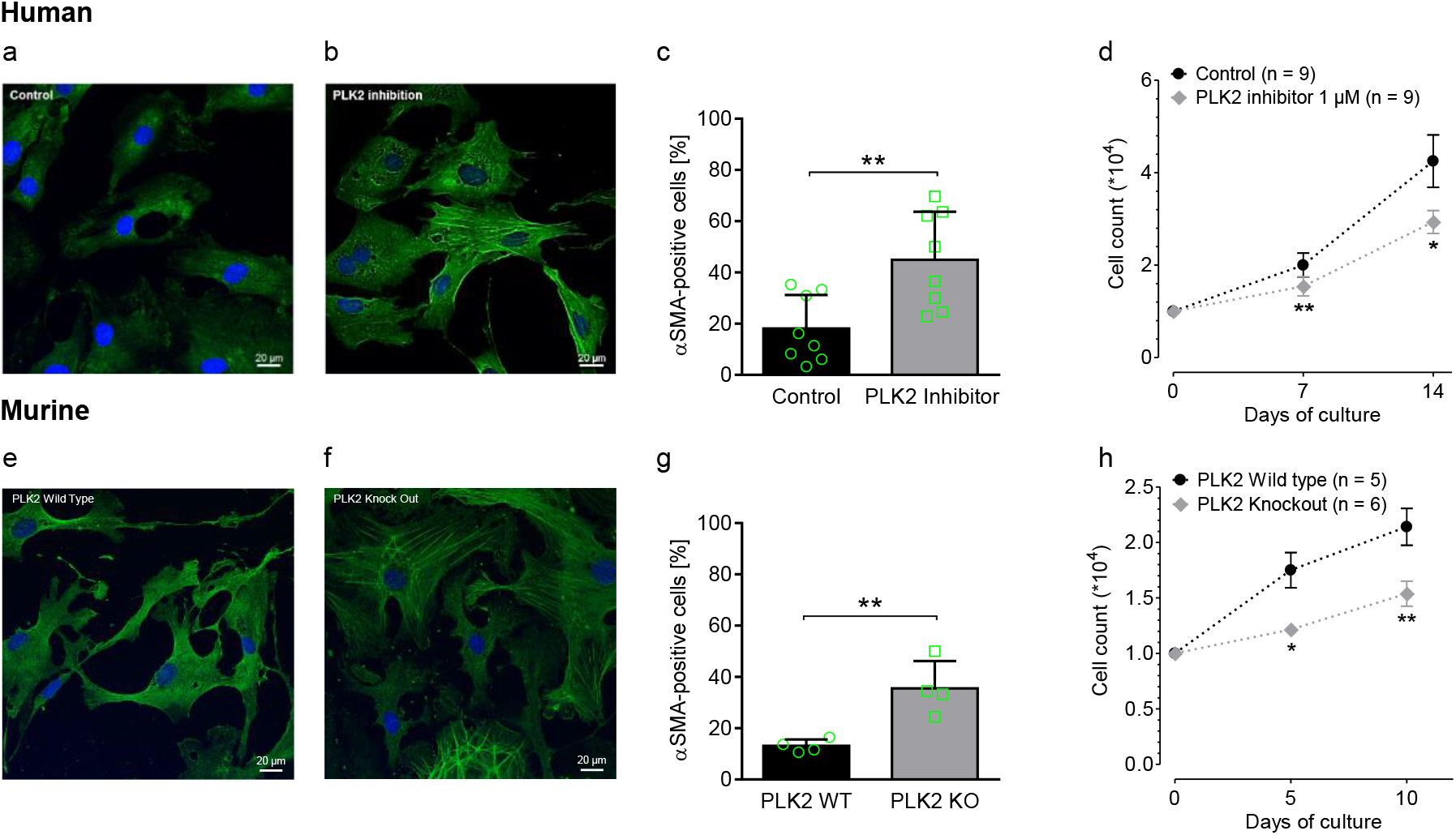
Inhibition and knockout of PLK2 induce myofibroblast differentiation and accordingly reduce fibroblast proliferation. (a – d): experiments performed with human atrial fibroblasts. **a)** and **b)** Immunofluorescence images stained for αSMA, the nuclei were stained with DAPI (blue). **c)** Quantification of immunostaining experiments for αSMA protein abundance dependent on PLK2 inhibition (n = 6 per group). Primary SR fibroblasts were incubated either with 1 μM TC-S 7005 or solvent control (1 μl DMSO/ ml medium) for 7 days. **d)** Proliferation curves of primary human SR fibroblasts. Cells were incubated either with 1 μM TC-S 7005 or solvent control (1 μl DMSO/ ml medium) (n = 9 per group). **(e – h): experiments performed with PLK2 wild type (WT) and knockout (KO) mouse fibroblasts. e)** and **f)** Immunofluorescence images stained for αSMA in PLK2 WT and KO fibroblasts, the nuclei were stained with DAPI (blue). **g)** Quantification of immunostaining experiments for αSMA protein abundance dependent on PLK2 expression (n = 4 mice per group). Primary PLK2 WT and KO fibroblasts were cultivated for 7 days. **h)** Proliferation curves of primary PLK2 WT and KO fibroblasts (n = 5 WT mice vs. 6 KO mice). ^*^: p < 0.05. ^**^: p < 0.01

### Inhibition and loss of PLK2 induce cellular senescence

Expression of senescence-associated β-galactosidase (SABG) is a hallmark of cellular senescence (Collado *et al*, 2007; Zhu *et al*, 2013). Hence, a screening assay of the SABG activity was performed with primary human SR fibroblasts treated with TC-S 7005 (Figure 4 a). Pharmacological PLK2 inhibition in human atrial fibroblasts resulted in significantly enhanced SABG activity compared to the vehicle-treated control group (Figure 4 b). This finding was confirmed in murine PLK2 WT and KO fibroblasts: PLK2 KO fibroblasts displayed higher basal SABG activity (Figure 4 c, p = 0.05) than WT.

**Figure 4:**
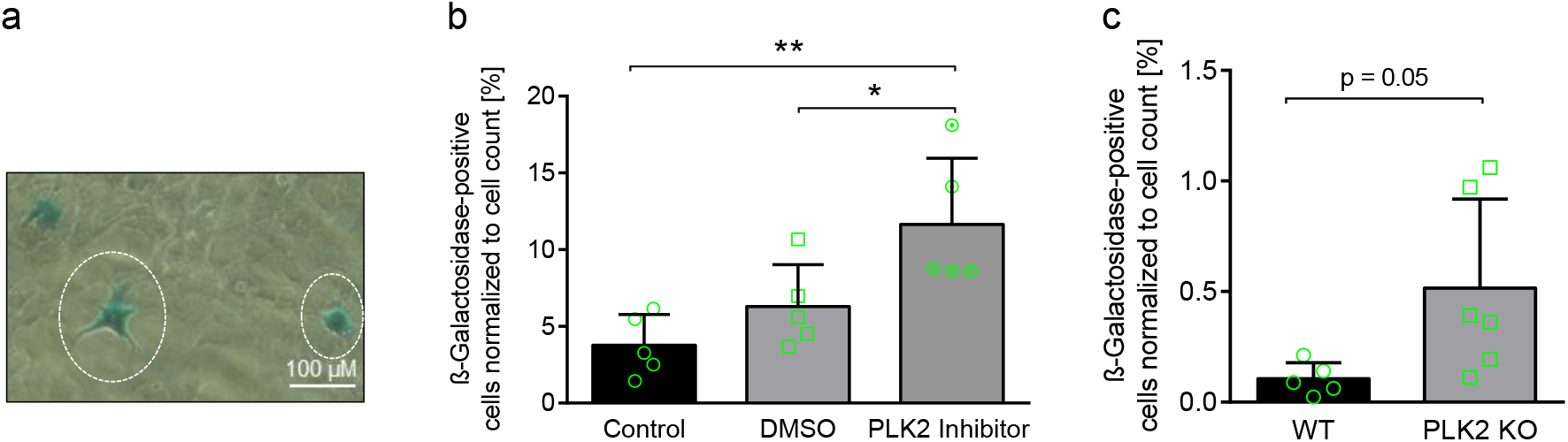
Loss of PLK2 expression and/ or function induces cellular senescence. **a)** Representative β-galactosidase-staining to detect cellular senescence. β-galactosidase-positive cells (senescent cells) are stained green. The scale bar equals 100 μm. **b** and **c)** Quantification of β-galactosidase-positive cells depending on PLK2 expression/ function. Values are depicted as percentage of the total cell count/ well. **b)** Proportion of senescent fibroblasts in human primary SR fibroblasts incubated with solvent control (1 μl DMSO/ ml medium) or 1 μM TC-S 7005 (PLK2 inhibitor) (n = 5 per group). **c)** Basal proportion of senescent cells in primary PLK2 WT and KO cardiac fibroblast culture (n = 5 WT mice vs. 6 KO mice). For statistical analysis a non-paired student‘s t-test was performed. For statistical comparison a one-way ANOVA with Newman-Keuls post test was performed. ^*^: p < 0.05. ^**^: p < 0.01

### Loss of PLK2 promotes *de novo* secretion of osteopontin

Based on the finding that PLK2 inhibition as well as KO initiated myofibroblast differentiation and senescence induction, we studied the secretory phenotype of PLK2 KO fibroblasts using mass spectrometry secretome analysis of the cell culture media (Coppé *et al*, 2008). We found particularly striking *de novo* secretion of OPN in PLK2 KO fibroblasts (Figure 5 a, supplemental table 4). Notably, OPN was also elevated in right atrial appendages from AF patients compared to SR (Figure 5 b). Consequently, we assessed levels of OPN in the peripheral blood of AF patients. To test if the severity of fibrosis in AF patients is directly correlated with serum OPN levels we compared AF patients with and without low voltage zones in the left atrium found during their electrophysiological study. Low voltage zones constitute the electrophysiological surrogate of atrial fibrosis (Verma *et al*, 2005). Both AF groups (without and with fibrosis) displayed elevated OPN levels in the peripheral blood (+1.7-fold in AF without fibrosis) compared to the SR group. However, OPN was 55% further elevated in AF patients with fibrosis compared to the non-fibrosis AF group (Figure 5 c). These results suggest fibrosis severity as a surrogate for fibroblast dysfunction correlating with peripheral blood OPN.

**Figure 5:**
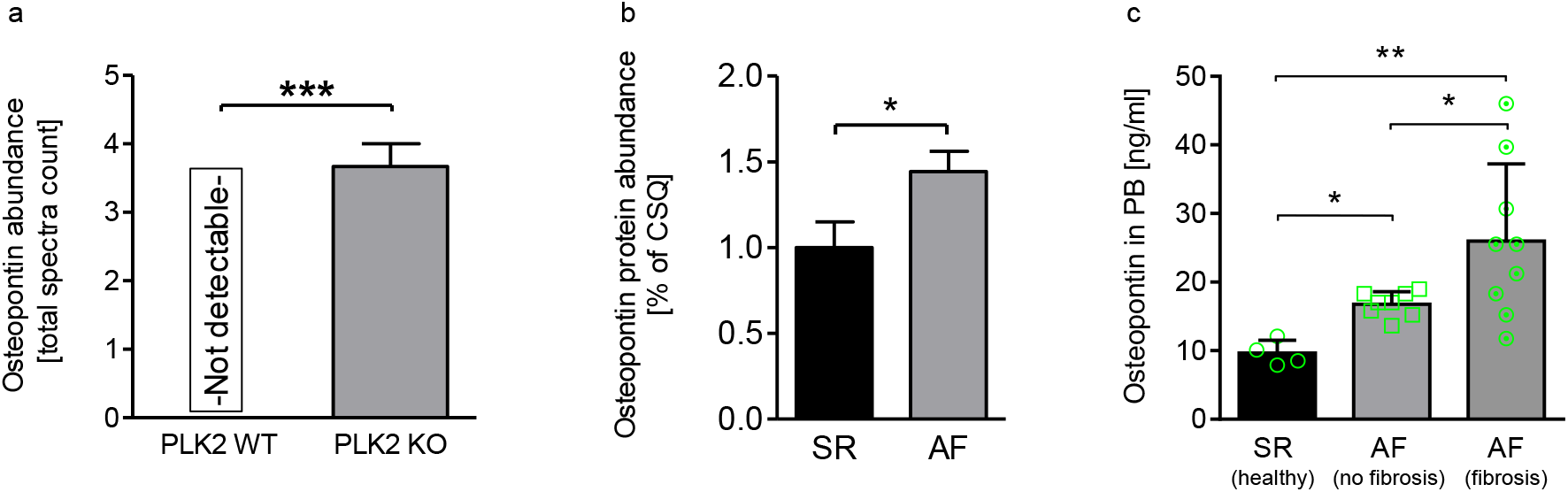
PLK2 mediates the expression of osteopontin. **a)** Osteopontin protein abundance in PLK2 WT or KO fibroblast culture medium analyzeded by mass spectrometry. PLK2 KO fibroblasts secrete osteopontin *de novo* (n = 3 mice per group). **b)** Quantification of western blots for osteopontin protein abundance in SR and AF right atrial tissue lysates (n = 10 per group). **c)** Osteopontin concentration in patients’ peripheral blood (PB). Osteopontin concentration was measured with ELISA (n_SR(healthy)_ = 4, n_AF(no fibrosis)_ = 8, n_AF(fibrosis)_ = 9). For statistical comparison in c) a one-way ANOVA with Newman-Keuls post test was performed. ^*^: p < 0.05. ^**^: p < 0.01. ^***^: p < 0.001

### Loss of PLK2 leads to exaggerated interstitial fibrosis

Since PLK2 knockout fibroblasts displayed significantly enhanced myofibroblast differentiation, we investigated the presence of interstitial fibrosis. Masson Trichrome staining of 4 months old PLK2 wildtype and knockout heart sections demarked vast interstitial fibrosis areas, especially in the left ventricle of PLK2 knockout animals compared to their wild type littermates which only displayed physiological septal collagen enrichment (Figure 6). Additional Sirius red staining of 8 months old PLK2 wildtype and knockout heart sections is displayed in Supplemental Figure 1.

**Figure 6:**
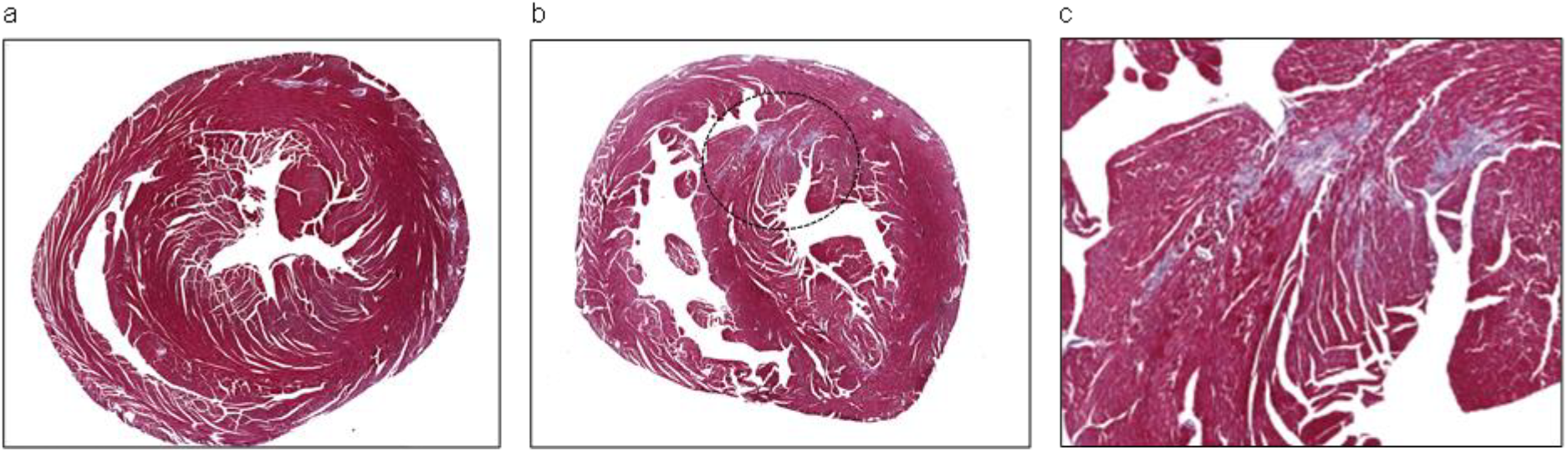
PLK2 knockout mice develop vast interstitial fibrosis compared to their wild type littermates. Paraffin sections of murine hearts. Collagen was stained intensively blue. The displayed area is located mid-ventricular (halfway between the cardiac valves and the apex cordis). **a)** PLK2 wildtype sample displayed as overview. **b)** PLK2 knockout sample displayed as overview. **c)** Magnification of **b** displaying an interstitial fibrosis area.

### Loss of PLK2 activates the p42/44 MAPK pathway *via* RasGRF2 to increase OPN expression

To propose a mechanistic link between lower PLK2 expression and higher OPN secretion of cardiac fibroblasts, we focused on the Ras - p42/44 MAPK pathway which was shown to act downstream of PLK2 as well as upstream of OPN (Hickey *et al*, 2005; El-Tanani *et al*, 2006). Indeed, inhibition of PLK2 with TC-S 7005 for 72h resulted in 2-fold higher expression of RasGRF2 in cardiac fibroblasts indicating that RasGRF2 protein abundance is regulated by PLK2 (Figure 7 a). We also found higher abundance of p42/44 MAPK (ERK1/2) in cardiac fibroblasts after TC-S 7005 pretreatment, which acts downstream of RasGRF2 and the Ras pathway (Thomas *et al*, 1992; Lee *et al*, 2011b, 2011a) (Figure 7 b, c). Several studies have shown that the Ras-pathway and p42/44 MAPK are mediators of OPN transcription (Hickey *et al*, 2005; El-Tanani *et al*, 2006; Xie *et al*, 2004). Accordingly, after inhibition of PLK2 we found higher OPN expression (p = 0.05) in fibroblasts (Figure 7 d). Representative western blots are provided in Figure 7 e – g. Thus, we provide evidence that reduced PLK2 expression in AF augments RasGRF2 and enhances expression of p42/44 MAPK which may stimulate the transcription of OPN and its secretion.

**Figure 7:**
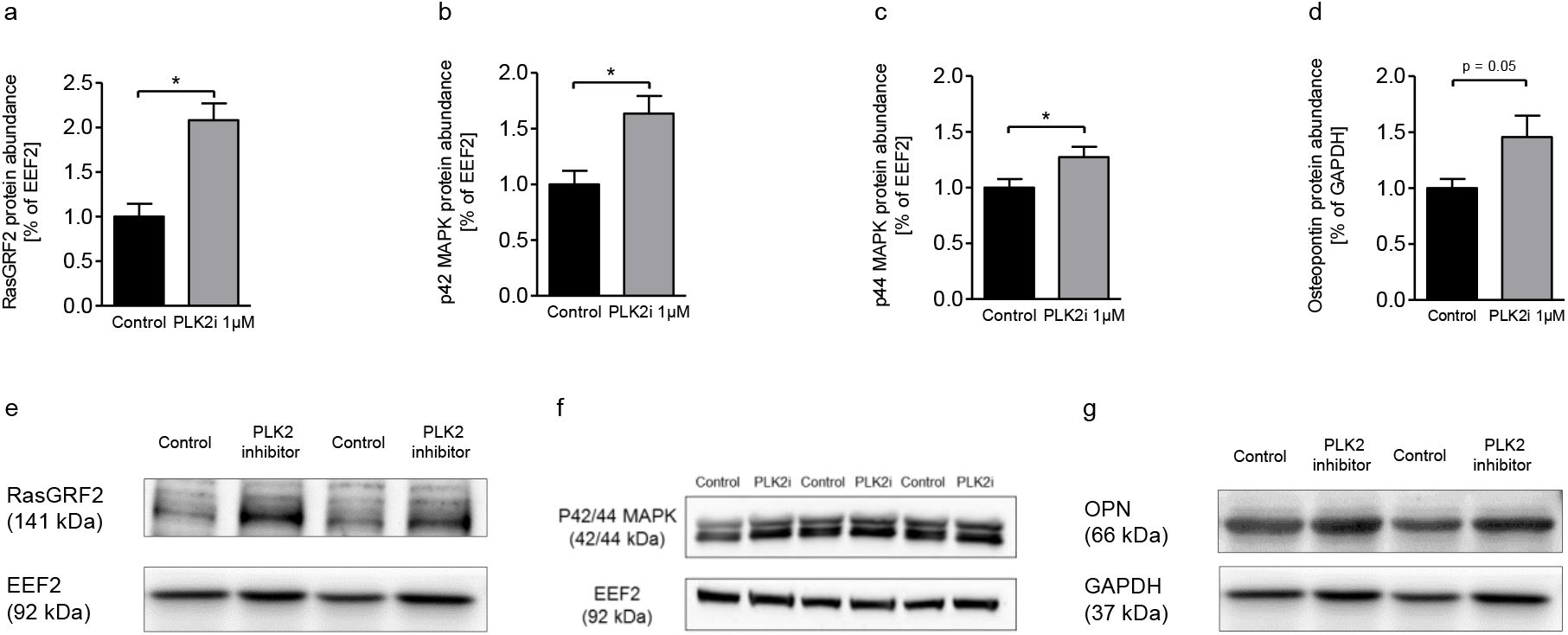
PLK2 regulates signal transduction and cytokines that induce inflammation. **a)** Quantification of RasGRF2 protein abundance in human ventricular fibroblasts treated with solvent control (DMSO 1 μl/ ml medium) or 1 μM TC-S 7005 (PLK2i) analyzed by western blot (n = 3 biological replicates per group). **b)** Quantification of p42 MAPK protein abundance in human ventricular fibroblasts treated with solvent control (DMSO 1 μl/ ml medium) or 1 μM TC-S 7005 (PLK2i) analyzed by western blot (n = 3 biological replicates per group). **c)** Quantification of p44 MAPK protein abundance in human ventricular fibroblasts treated with solvent control (DMSO 1 μl/ ml medium) or 1 μM TC-S 7005 (PLK2i) analyzed by western blot (n = 3 biological replicates per group). **d)** Quantification of OPN protein abundance in human ventricular fibroblasts treated with solvent control (DMSO 1 μl/ ml medium) or 1 μM TC-S 7005 (PLK2i) analyzed by western blot (n = 3 biological replicates per group). **e, f, g)** Representative western blots for RasGRF2, p42/44 MAPK and OPN. ^*^: p < 0.05.

## Discussion

This study demonstrates that dysregulated PLK2 contributes to fibroblast activation, subsequent myofibroblast transition and enhanced secretion of OPN. Moreover, PLK2 expression is sensitive to established AF-dependent stress conditions such as hypoxia. Thus, its druggable downstream effectors (Ras/ MAPK/ OPN) may constitute valuable targets for antifibrotic and/or anti-inflammatory interventions to abolish pathological atrial fibrosis in patients with AF (Figure 8).

**Figure 8:**
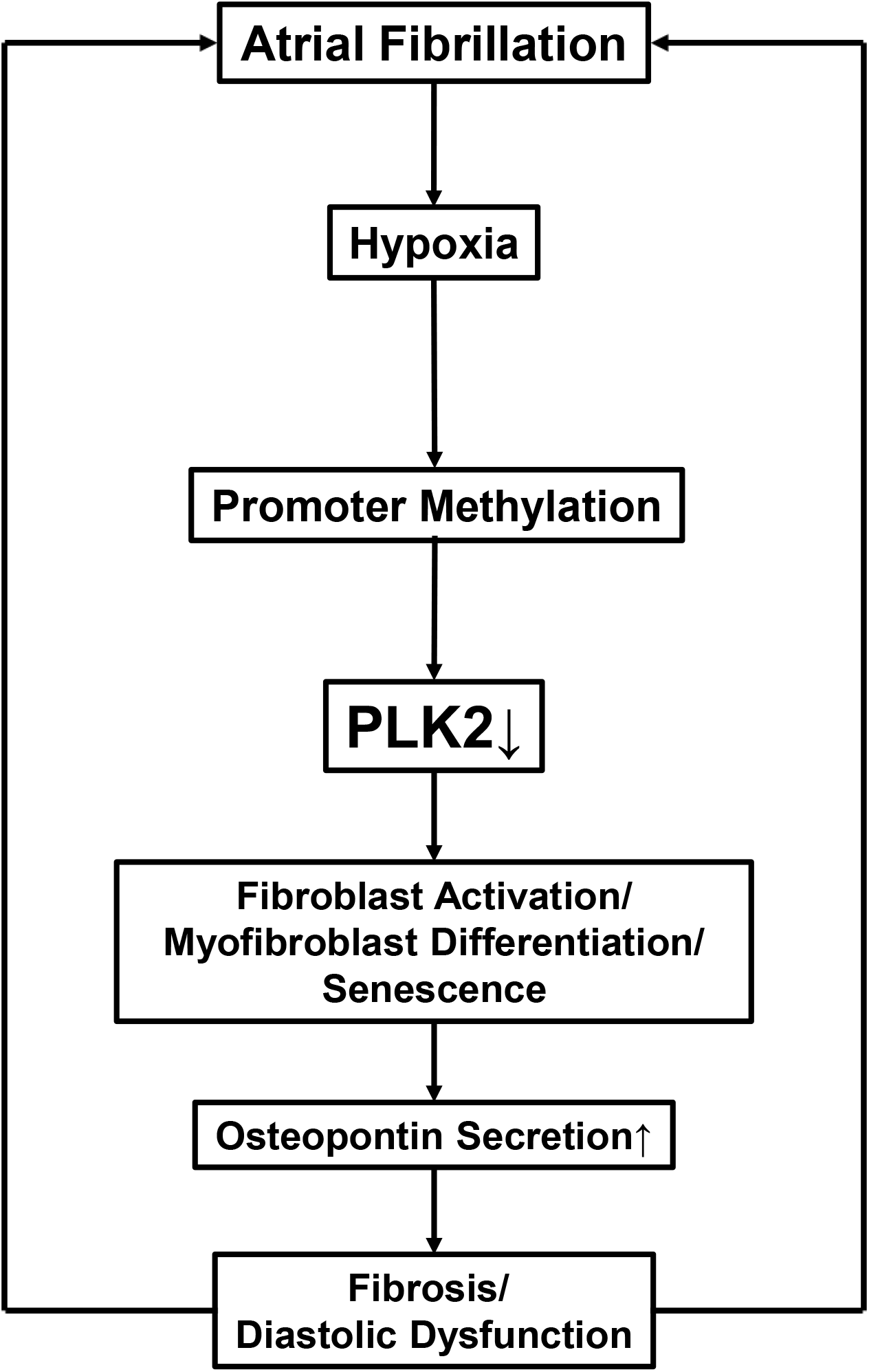
Proposed model for AF-induced PLK2 downregulation and concomitant downstream events. Prolonged tissue hypoxia is present in AF and induces *PLK2* promoter methylation. Downstream of lower PLK2 expression (mRNA and protein) fibroblasts are activated and differentiate into myofibroblasts or go into cell senescence. Those metabolically active fibroblast populations *de novo* secrete the inflammation mediator osteopontin which causes additional fibrosis and worsens diastolic dysfunction.

Hypoxia has already been linked to global DNA methylation, contributing to fibrosis development in the lung by inducing myofibroblast differentiation and the expression of fibrosis markers (Robinson *et al*, 2012). We observed pronounced PLK2 downregulation already after 24h of hypoxia treatment (1% O_2_) (Figure 2 c) but increased *PLK2* promoter methylation was not detectable until 4 days of hypoxia treatment. This is in line with a recent publication which identified 4 days of hypoxia as minimum to reach the threshold for detection of DNA methylation in human fibroblasts (Robinson *et al*, 2012). Our finding of aberrant methylation in AF patients is of clinical interest, since DNA methylation is reversible (Plumb *et al*, 2000): In a mouse model of kidney fibrosis, demethylation of the *RASAL1* gene by application of the DNA methyl transferase inhibitor 5-azacytidine significantly attenuated pathological fibrosis (Bechtel *et al*, 2010). For PLK2, a similar approach could be feasible to counteract fibroblast activation and fibrotic remodeling in AF.

Furthermore, we provide a link between PLK2 and the fate of cardiac fibroblasts. We demonstrate that PLK2 inhibition or KO induce myofibroblast differentiation and consequently reduce cell proliferation in cardiac fibroblasts. This finding fits well to a recent study showing PLK2 as a key regulator for proliferation and differentiation in cardiac progenitor cells (Mochizuki *et al*, 2017). Moreover, primary human cardiac fibroblasts from AF patients were shown to differentiate into myofibroblasts to a greater extent and to proliferate less compared to SR controls (Poulet *et al*, 2016). Activated myofibroblasts are highly active secretory cells that substantially contribute to fibrosis in the diseased heart (Tallquist & Molkentin, 2017). Accordingly, we demonstrated vast interstitial fibrosis areas in the PLK2 knockout mice by histological analysis. Fibrosis is considered then to worsen cardiac performance by increasing wall stiffness leading to reduced diastolic filling, increased heart rate and consequently increased oxygen consumption. Moreover, fibrosis underlies the arrhythmogenic substrate fostering cardiac arrhythmia like AF (Figure 8). Our data suggest that reduced PLK2 expression in AF is a molecular trigger for enhanced myofibroblast differentiation and lower proliferation in AF.

Importantly, beside enhanced myofibroblast differentiation inhibition of PLK2 led to accelerated cellular senescence in primary human atrial fibroblasts. This finding is consistent with a study showing that transcriptional inhibition of PLK2 by miRNA-146a induced senescence in bone marrow cells (Deng *et al*, 2017). There is growing evidence that increased proportions of senescent fibroblasts can contribute to the disruption of regular tissue architecture (Krtolica *et al*, 2001; Parrinello *et al*, 2005). Moreover, cardiac senescence has been shown to play an important role in diabetic cardiomyopathy and fibrosis in a mouse model (Gu *et al*, 2018). In this scenario, we propose that PKL2 is a novel regulator of cardiac fibroblast senescence, which may additionally contribute to the pathogeneses of AF patients.

Myofibroblasts and senescent fibroblasts are known to secrete a plethora of extracellular matrix proteins and inflammatory cytokines (Baum & Duffy, 2011; Childs *et al*, 2015). In PLK2 KO fibroblast culture media, discovery proteomics revealed *de novo* secretion of the inflammatory cytokine-like protein OPN which has been shown to induce inflammation, arteriosclerosis and cardiomyopathy (Zhao *et al*, 2016). In AF patients, OPN was elevated on tissue-level in right atria as well as systemically in the peripheral blood in comparison to SR controls. OPN levels in the peripheral blood of patients have been investigated as prognostic biomarker to predict the AF recurrence after cryoballoon catheter ablation (Güneş *et al*, 2017). Additionally, high OPN plasma levels correlate with ventricular tachycardia and fibrillation in heart failure patients (Francia *et al*, 2014). Our data indicate that OPN is a PLK2-dependent systemic-inflammatory component of AF, which correlates with concomitant cardiac fibrosis.

Finally, we propose a mechanistic link between PLK2 suppression and enhanced OPN secretion. We found that RasGRF2 is downstream regulated by PLK2 and that inhibition of PLK2 leads to p42/44 MAPK (ERK1/2) accumulation in cardiac fibroblasts. PLK2 phosphorylates RasGRF1 leading to its proteasomal degradation and thereby inhibits the Ras pathway (Lee *et al*, 2011a). Moreover, the Ras-pathway and p42/44 MAPK have been shown to modulate OPN transcription (Hickey *et al*, 2005; El-Tanani *et al*, 2006; Xie *et al*, 2004). Reduced phosphorylation-dependent degradation of RasGRF2 by inhibition of PLK2 may well underlie our findings in the present study. RasGRF2 on the other hand is known to stimulate the Ras pathway resulting in higher expression of p42/44 MAPK which then may stimulate OPN expression and secretion. To study whether our findings reflect aberrant gene expression, RNA sequencing with subsequent bioinformatical analysis was performed in PLK2 WT and KO fibroblasts. Surprisingly, there were no significantly altered mRNA expression patterns between PLK2 WT and KO (except for missing PLK2 expression in the KO group) (Supplemental figure 2). This finding thus indicates that loss of PLK2 affects mainly protein expression on a post transcriptional level. Our hypothesis of OPN being the key mediator of PLK2-dependent inflammation and fibrosis in AF could be of particular clinical relevance because of existing and approved compounds counteracting OPN (Geschka *et al*, 2011), e.g. soluble guanylate cyclase stimulators like riociguat which ameliorate cardiac fibrosis and attenuate OPN release (Geschka *et al*, 2011; Sandner *et al*, 2017). These small molecules could be a starting point to modulate the PLK2-OPN axis in treatment and prevention of AF.

## Conclusions and Clinical Implications

AF, the most relevant cardiac arrhythmia in humans, is a complex, systemic inflammatory disorder with strikingly increasing numbers of afflicted patients. Yet, only little is known about the underlying molecular pathways. Our work demonstrates that the PLK2-OPN axis in atrial fibroblasts contributes to fibroblast dysfunction and pathological fibrosis in permanent AF. To our knowledge, this is the first study to propose a novel pathophysiological role for the family of polo-like kinases in AF pathophysiology. Here we provide a mechanistic link between reduced PLK2 expression in AF and concomitant OPN release. Our results suggest that modulation of PLK2 activity and/or inhibition of OPN release could ameliorate pathological remodeling in AF.

## Methods

### Human sample acquisition

All patients participating in our study gave written informed consent according to the Declaration of Helsinki and the study was approved by the institutional review committee (Official file numbers: EK 114082202, EK 465122013). Right atrial appendages were collected in collaboration with the Herzzentrum Dresden GmbH. Patients undergoing open heart surgery like bypass surgery or valve replacement donated their right atrial appendages that incurred as a result of the surgical procedure. The samples were put into 4° C cold isotonic transport solution and immediately processed for fibroblast isolation. Whenever excess material remained, the tissue was frozen and stored in liquid nitrogen. Peripheral blood samples from AF patients who would undergo ablation of pulmonary veins were collected prior to the intervention in EDTA-tubes. Low voltage zones were assessed by electrophysiological mapping of the left atrium and defined as bipolar voltage < 0.5 mV. Blood samples were kept at 4° C for a maximum of 1 h. The tubes were then centrifuged for 10 min at 1000 g at 4° C. Subsequently, the plasma was transferred carefully into 500 μl Eppendorf tubes and stored at -80° C until analysis. Corresponding patient data can be found in supplemental tables 1, 2 and 3.

### PLK2 WT and KO mice

PLK2 WT and KO mice are commercially available via The Jackson Laboratory (Bar Harbor, USA) (129S.B6N-*Plk2^tm1Elan^*/J, stock number: 017001 Plk2 KO). The mouse model has previously been used to study the alpha-synuclein phosphorylation at serine 129 in the central nervous system (Inglis *et al*, 2009). The animal study was approved by the institutional bioethics committee (T 2014/5; TVA 25/2017).

### Cell isolation and culture

Primary human right atrial fibroblasts were isolated via outgrowth method from right atrial appendage biopsies as published previously(Poulet *et al*, 2016). Human ventricular fibroblasts were purchased from abm Inc. (Richmond, Canada) and adapted to the culture conditions of the primary cells. Primary murine cardiac fibroblasts were isolated enzymatically via Langendorff-perfusion(El-Armouche *et al*, 2008). The supernatant was centrifuged for 1 min at 350 g to remove cardiomyocytes and debris. The resulting supernatant was centrifuged a second time for 1 min at 750 g to sediment fibroblasts. Cells were cultured in Dulbecco modified eagle medium (Life Technologies, Carlsbad CA, USA) supplemented with 10 % fetal calf serum (FCS, Life Technologies, Carlsbad CA, USA) and 1

% penicillin/ streptomycin (Life Technologies, Carlsbad CA, USA). The cell culture medium was changed every other day. Equal volumes of the specific PLK2 inhibitor TC-S 7005 (Tocris Bioscience, Bristol, UK) and the solvent control (DMSO) were added with the medium change. Cells were kept in a humidified surrounding at 37°C and 5 % CO_2_. In order to generate hypoxic conditions (1 % O_2_) cells were cultured in a hypoxia incubator chamber (Coy Laboratory Products Inc, Grass Lake, USA) for 24 up to 72 h.

### Immunocytofluorescence

In order to perform immunocytoflorescence experiments, fibroblasts were seeded on 1 cm glass cover slips and grown for 7±1 days until they reached approximately 80 % of optical confluence. Cells were then washed with cold PBS, fixed in 4 % PFA for 15 min at RT, washed and subsequently permeabilised using Triton X. After blocking with FCS the cover slips were incubated with a mixture of primary antibody against αSMA (Sigma-Aldrich, St. Louis, Missouri, USA) and 4′,6-Diamidin-2-phenylindol (DAPI) (1:200; A5228, Sigma-Aldrich, St. Louis, Missouri, USA) for 1 h in a dark, humidified surrounding at RT. After washing, the secondary antibody Alexa-Fluor 448 (Abcam, Cambridge, UK) was

applied for 1 h in a dark, humidified surrounding at RT. The samples were kept in the dark until fluorescence images were obtained with a Zeiss LSM-510 confocal microscope.

### Histological analysis

For histological analysis, a mid-ventricular section of the heart was excised and fixed in 4% PFA for 24 h before dehydration using an ethanol gradient. Subsequently the samples were embedded in paraffin, sectioned at 3 μm thickness and stained with either Masson Trichrome or Sirius red (Sigma-Aldrich, St. Louis, Missouri, USA). Images were acquired using the Keyence BZ-X710 All-in-One Fluorescence Microscope (Keyence Corporation of America, Itasca, USA).

### β-galactosidase staining assay for senescence detection

Fibroblasts were plated on 6-well plates at densities of 2.5^*^10^4^ cells/ well and kept in culture for 7 days. Subsequently, cells were fixed with 4 % formaldehyde for 15 min at room temperature (RT) and the senescence staining was done. A standard senescence detection kit (senescence associated β-galactosidase (SABG) staining) was used according to the manufacturer’s instructions (Abcam, Cambridge, UK). SABG-positive cells were stained green and considered senescent (Figure 4 a).

### Quantitative real-time PCR (QPCR)

SYBR green (Bio-Rad Laboratories GmbH, Munich, Germany) real-time PCR was used to measure the gene expression of PLK2 *in vitro*. Specific primers for PLK2 were purchased from Bio-Rad (Bio-Rad Laboratories GmbH, Munich, Germany). GAPDH was used as housekeeping gene. For RNA isolation and subsequent cDNA synthesis the PeqLab total RNA mini and PeqGold cDNA synthesis kits (Peqlab Biotechnologie GmbH, Erlangen, Germany) were used according to the manufacturer’s instructions. Optional on-column DNA digestion was performed, in order to remove residual contaminating genomic DNA. PCR runs were performed in a CFX96 Touch Deep Well Real-Time PCR detection system (Bio-Rad Laboratories GmbH, Munich, Germany). Samples were amplified in duplicates or triplicates as indicated in the results part. For data analysis the CFX manager software (Bio-Rad Laboratories GmbH, Munich, Germany) was used. Relative gene expression was calculated to housekeeping gene.

### Methylation-specific PCR

The methylation-specific PCR was performed as published by Syed *et al*., 2006and Robinson *et al.,* 2017(Syed *et al*, 2006; Benetatos *et al*, 2011). Genomic DNA (gDNA)was isolated using the PureLink Genomic DNA Extraction kit (Thermo Fisher Scientific, Waltham, Massachusetts, USA). Purified gDNA was subsequently bisulfite-converted using the EZ DNA starter kit according to the manufacturer’s instructions. The following PCR protocol was designed according to the suggestions of ZYMO Research. For unmethylated samples 36 cycles were run and for detection of DNA methylation 38 runs, respectively. For electrophoresis, the PCR products were then applied to a 2 % agarose gel containing HD green (INTAS Science Imaging Instruments GmbH, Göttingen, Germany). Visualization of gel bands was achieved with a Fusion FX (Peqlab Biotechnologie GmbH, Erlangen, Germany) development device. The following primers as published by Syed *et al.*(Syed *et al*, 2006)were used for methylation-specific PCR:

PLK2 unmethylated for.: 5′-CACCCCACAACCAACCAAACACACA-3′

PLK2 unmethylated rev.: 5′-GGATGGTTTTGAAGGTTTTTTTGTGGTT-3′ (product = 142 bp) PLK2 methylated for.: 5′-CCCACGACCGACCGAACGCGCG-3′

PLK2 methylated rev.: 5′-ACGGTTTTGAAGGTTTTTTCGCGGTC-3′ (product = 137 bp)

### SDS-PAGE, Western Blotting and Immunodetection

Protein was extracted from whole heart tissue and cells for western blot analysis using RIPA buffer (30 mM Tris, 0.5 mM EDTA, 150 mM NaCl, 1% NP-40, 0.1 % SDS) supplemented with 10 % protease and phosphatase inhibitors. Protein concentration was determined using a BCA kit (Pierce Biotechnology, Waltham, Massachusetts, USA). For SDS-PAGE, 10 % polyacrylamide gels were used. 30 μg of protein were loaded into each lane of the gels. Proteins were subsequently transferred to a 0.45 μm nitrocellulose membrane. Equal loading was ensured by ponceau red staining (Sigma-Aldrich, St. Louis, Missouri, USA) before blocking in 5 % milk. Membranes were incubated with primary antibodies overnight at 4°C under constant gentle shaking. After several washing steps, secondary antibodies (anti mouse or anti rabbit) were applied for 1 h at RT. After final washing membranes were incubated with ECL development solution (Thermo Fisher Scientific, Waltham, Massachusetts, USA) and developed in a Fusion FX device (Vilber Lourmat Deutschland GmbH, Eberhardzell, Germany). Depending on the molecular weight of the proteins of interest either Glyceraldehyde 3-phosphate dehydrogenase (GAPDH), calsequestrin (CSQ) or Eukaryotic elongation factor 2 (EEF2) were used as housekeeping proteins.

**Table 1:** Western blot antibodies

| Antibody | Dilution | Origin | Supplier |
| --- | --- | --- | --- |
| GAPDH | 1 : 1000 | Mouse | Santa-Cruz, sc-365062 |
| EEF2 | 1 : 50000 | Rabbit | Abcam, ab40812 |
| PLK2 | 1 : 1000 | Rabbit | Cell Signaling, #14812 |
| αSMA | 1 : 200 | Mouse | Sigma-Aldrich, A5228 |
| RasGRF2 | 1 : 1000 | Rabbit | Sigma-Aldrich, SAB2700396 |
| p42/44 MAPK (ERK1/2) | 1 : 1000 | Rabbit | Cell Signaling, 9102 |
| OPN | 1 : 1000 | Rabbit | Abcam, ab8448 |

### Secretome analysis using LC-MS/MS(Yin *et al*, 2013; Abonnenc *et al*, 2013)

The fibroblasts were isolated from PLK2 wild type and knock out mouse and cultured in serum-free medium for 72 hours. Conditioned media were collected and concentrated using 3kD molecular weight cut off spin columns (Amicon, Millipore, Bedford, USA) and washed 5x with 25mM ammonium bicarbonate. The samples were denatured using 8M urea/ 2M thiourea and reduced by 10mM DTT. After alkylated with 50mM iodoacetamide, the samples were digested using trypsin (enzyme:protein=1:20) overnight. Digested peptides were purified using C18 spin plate (Harvard Apparatus, Holliston, USA). The eluted peptides were resuspend in LC solution (2% acetonitrile, 0.05% TFA) and 1ug was injected and separated by reverse phase nano-flow HPLC (Dionex UltiMate 3000 RSLCnano, Acclaim PepMap100 C18 column, 75um × 50cm, Thermo Fisher Scientific, Waltham, Massachusetts, USA). The nanoflow mobile phases consisted of HPLC grade water containing 0.1% formic acid (mobile phase A) and acetonitrile/HPLC grade water (80:20, v:v) containing 0.1% formic acid (mobile phase B). The following gradient was run at 250nL/min: 0-10 min, 4-10% B; 10-75 min, 10-30% B; 75-80 min, 30-40% B; 80-85 min, 40-99% B; 85-89.8 min, 99% B; 89.8-90 min, 99-4% B; 90-120 min, 4%B. The nano column was coupled to a nanospray source (Picoview, New Objective, Woburn, USA). Spectra were collected from a Q Exactive HF (Thermo Fisher Scientific, Waltham, Massachusetts, USA) in positive ion mode using Full MS resolution 60,000 (at 200 m/z), scan range 350 to 1600 m/z. Data-dependent MS/MS was performed using higher-energy collisional dissociation (HCD) fragmentation on the 15 most intense ions with resolution 15,000 and dynamic exclusion enabled. Raw files were searched against UniProt/SwissProt Mouse and Bovine database (version 2016_02, 22763 protein entries) using Proteome Discoverer 1.4.1.14. The mass tolerance was set at 10 ppm for the precursor ions and at 20 mmu for fragment ions. Carbamidomethylation of cysteine was set as a fixed modification, oxidation of methionine, proline and lysine was set as variable modifications. Two missed cleavages were allowed. Search result files were loaded into Scaffold software (version 4.3.2) and validated with the following filter: peptide probability > 95% and protein probability > 99% with minimum 2 peptides. Total spectra count was used as quantitative value.

### RNA sequencing and bioinformatics

The RNA sequencing and subsequent bioinformatical analysis was done as published by Schott *et al*(Schott *et al*, 2017). For Whole-Transcriptome Sequencing (RNA-Seq), 1.5× 10^6^ murine cardiac PLK2 WT and KO fibroblasts were seeded in T-25 flasks and cultured until optical confluency. Subsequently, cells were harvested for RNA extraction (as described above). For library preparation the TruSeq Stranded Total RNA Library Prep Kit (Illumina, San Diego, USA) was used according to the manufacturer’s instructions, starting with 1 μg total RNA. All barcoded libraries were pooled and sequenced 2×75bp paired-end on an Illumina NextSeq500 platform to obtain a minimum of 10 Mio reads per sample. Raw reads were converted from bcl to fastq format using bcl2fastq, quality trimmed with trimmomatic(Bolger *et al*, 2014) and then aligned against the Ensembl 92 mouse genome using STAR(Dobin *et al*, 2013) in a 2-pass mapping mode. Read counts of all annotated genes were extracted from the alignments using feature Counts method of the Rsubread package(Liao *et al*, 2013) and genes with 0 counts for all samples were discarded. DESeq2(Love *et al*, 2014) was used to find differentially expressed genes using standard parameters. Only genes with multiple testing adjusted p-values < 0.05 were considered significant. Clustering was done with Euclidean distance and complete linkage using regularized-logarithm transformation (rlog) of TPM (Transcripts Per Kilobase Million) expression values. Heatmaps were plotted using the R package ComplexHeatmap(Gu *et al*, 2016) and hclust of the R stats package. Principal components analysis was done using the R stats package(R Core Team, 2014).

### Human OPN assay

The human OPN ELISA kit was purchased from Abcam (Abcam, Cambridge, UK) and used according to the manufacturer’s instructions to detect the concentration of OPN in the peripheral blood of control SR controls and AF patients.

### Statistical analysis

Results are presented as mean ± SEM. For statistical analysis and graphic representation of the data, Graph Pad Prism software v.5 (GraphPad Software, San Diego, USA) was used. For comparisons between two groups, student’s t-test was used with Welsh’s correction if appropriate. When comparing three groups, a one-way ANOVA with Newman-Keuls posttest was performed. To compare the presence or absence of promoter methylation (Figure 2) Fisher’s exact test was used. P-values < 0.05 were considered statistically significant and indicated with asterisks (^*^) in the corresponding figures (^*^*p* < *0.05*; ^**^*p* < *0.01*; ^***^*p* < *0.001*).

## Acknowledgments

We thank the “Förderkreis der Dresdner Herz-Kreislauf-Tage” for supporting this study with the scientific research grant “Forschungspreis der Dresdner Herz-Kreislauf-Tage”. We are grateful to Annett Opitz and Romy Kempe for expert technical assistance. We want to thank Charlotte Schaeffer, Pia Schlinkert and Björn Binnewerg for support in sample generation and PCR analysis. We are grateful to Stephanie M. Schacht for expert assistance in animal keeping and handling. We are indebted to Prof. Dr. Samuel Sossalla (Regensburg university hospital, Germany) for providing RNA samples from human cardiomyocytes.

## Author contributions

S.R.K. designed and performed the experiments, acquired funding, analyzed the data, prepared the figures and wrote the paper. K.S. designed and performed experiments and analyzed the data. T.K. contributed to the manuscript preparation. S.M.R. supervised the animal work. C.P. provided clinical data and blood samples. S.M.T. provided primary human heart tissue samples for subsequent cell cultures. S.R.J. provided the PLK2 KO animals and expert knowledge. M. M. designed experiments and provided support for the mass-spectrometry. X.Y. performed experiments and provided support for the analysis of mass-spectrometry data. J.D.K., P.W. and K.Gr. designed and performed experiments and contributed to manuscript preparation. N.H. and J-H.K. contributed to manuscript preparation. U.R., M.W., S.K. and K.G. supervised the experiments and manuscript preparation. S.W. designed experiments and contributed to manuscript preparation. A.E.A. supervised the work, acquired funding and wrote the paper.

## Competing interests

All authors concur with the submission of the manuscript and none of the data have been previously reported or are under consideration for publication elsewhere. Parts of S.R.K.’s doctoral thesis are included in this manuscript. There are no competing interests or conflicts of interests to declare.

## The Paper Explained

### Problem

Although atrial fibrillation and subsequent fibrotic remodeling of the heart affect millions of patients worldwide, the underlying molecular mechanisms through which atrial fibrillation leads to heart remodeling and fibrosis in patients are incompletely understood. Consequently, there is a need to define targets for therapies that limit the debilitating consequences of atrial fibrillation.

### Results

We identified PLK 2 as an epigenetically regulated kinase involved in the pathophysiology of fibrosis in atrial fibrillation. In atrial tissue samples from patients with chronic atrial fibrillation, PLK 2 was nearly 50 % downregulated by hypoxia-induced promoter methylation. *In vitro*, loss of PLK 2 led to strikingly reduced fibroblast proliferation, increased myofibroblast differentiation and enhanced senescence induction. Additionally, we found that genetic knockout of PLK2 resulted in *de novo* secretion of the inflammatory cytokine-like protein osteopontin. Accordingly, we measured *ex vivo* elevated osteopontin in both heart tissue and the peripheral blood of atrial fibrillation patients and healthy sinus rhythm controls. We identified the Ras-ERK1/2 signaling pathway to be the potential link between reduced expression of PLK2 and elevated osteopontin secretion.

### Impact

Our findings reveal PLK 2 to be a regulator of cardiac fibroblast function, inflammation and fibrosis development. Our results strengthen the current hypothesis that atrial fibrillation is not only an ion channel disease but a complex (systemic) inflammatory disorder. Restoration of physiological PLK2 expression or blockade of osteopontin release may constitute valuable new drug targets for the prevention and treatment of cardiac fibrosis in atrial fibrillation.

## Supplementary Material

**Supplemental Figure 1:**
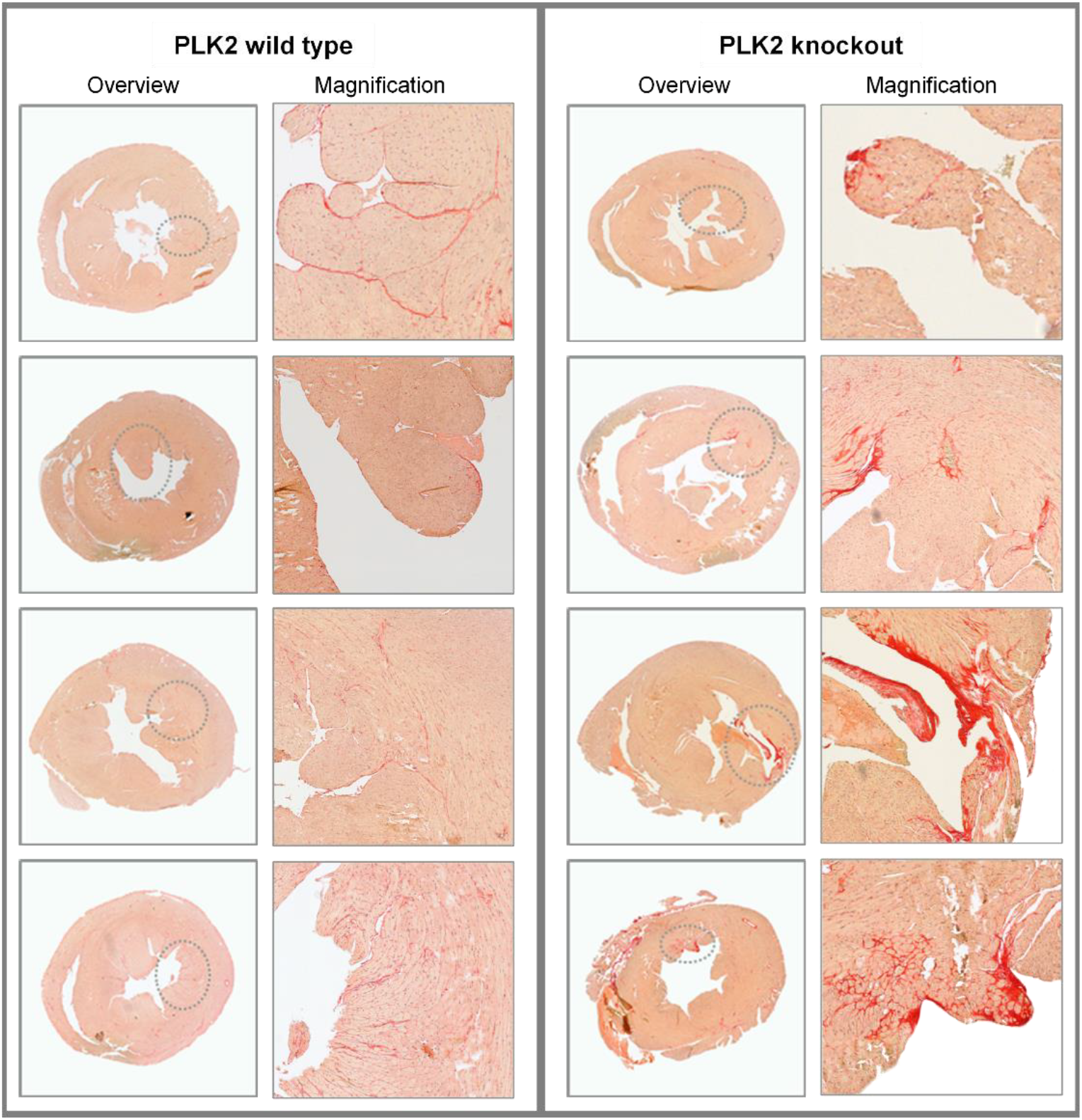
Sirius red staining of histological sections of PLK2 wild type and knockout hearts. Paraffin sections of murine hearts. Collagen was stained intensively red. The area displayed is located mid-ventricular (halfway between the cardiac valves and the apex cordis). **Left panel)** PLK2 wildtype samples (n = 4) displayed as an overview and with a corresponding magnified area. **Right panel)** PLK2 knockout samples (n = 4) displayed as an overview and with a corresponding magnified area.

**Supplemental Figure 2:**
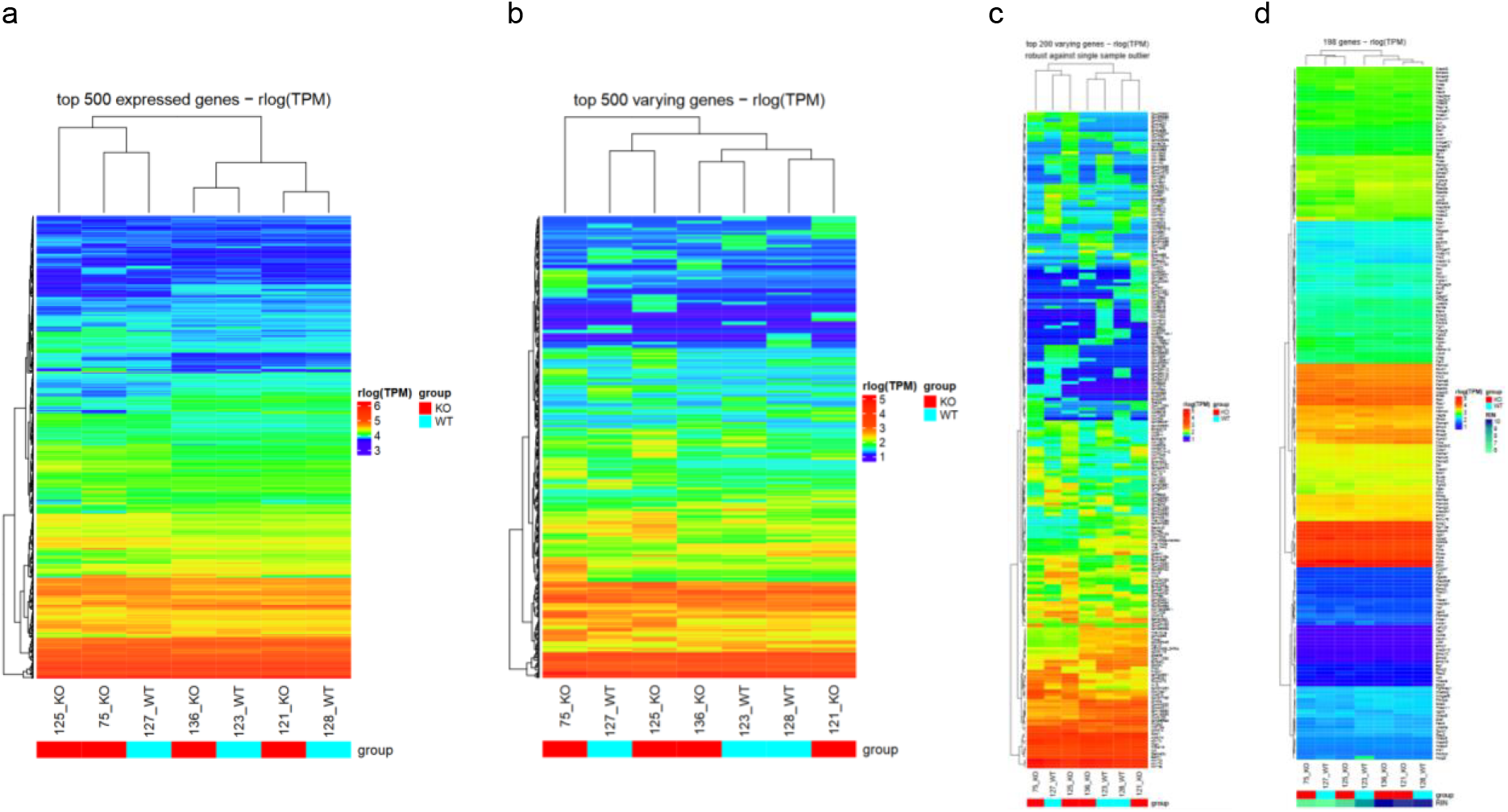
Clustering of PLK2 WT and KO mouse fibroblast RNA sequencing. Analysis of the RNA sequencing revealed no significantly regulated genes between PLK2 WT and KO except for missing PLK2 expression in the KO group indicating that KO of PLK2 alters solely post transcriptional signaling. **a)** Heat map depicting the top 500 expressed genes in PLK2 WT and KO fibroblasts. **b)** Heat map depicting the top 500 varying genes between PLK2 WT and KO. **c)** Heat map depicting the top 200 varying genes between PLK2 WT and KO. **d)** Heat map depicting genes involved in Ras and SMAD signaling.

**Supplemental Table 1:** Patient data for the human osteopontin ELISA

|  | Healthy (n=4) | AF no fibrosis (n=8) | AF with fibrosis (n=9) |
| --- | --- | --- | --- |
| **Average age [mean, years]** | 54 | 71,3 | 71,4 |
| **Gender** |  |  |  |
| Male | 2 | 5 | 5 |
| Female | 2 | 3 | 4 |
| **Disease** |  |  |  |
| Hypertension | 1 | 8 | 9 |
| Diabetes mellitus | 0 | 4 | 3 |
| Hyperlipidemia | 0 | 5 | 5 |
| Chronic kidney disease (GFR 40 - 90) | 0 | 2 | 3 |
| Chronic lung disease | 0 | 1 | 2 |
| Thyroid disease | 0 | 0 | 2 |
| Adipositas | 0 | 1 | 0 |
| Current smoking | 0 | 1 | 1 |
| **Atrial fibrillation characteristics** |  |  |  |
| Persistent AF | 0 | 4 | 7 |
| Paroxysmal AF | 0 | 3 | 1 |
| Atrial flutter | 0 | 1 | 1 |
| **Drugs** |  |  |  |
| ACE inhibitors | 1 | 0 | 6 |
| AT1 receptor blockers | 0 | 5 | 1 |
| β-AR blockers | 0 | 7 | 9 |
| Calcium channel blockers (nifedipine) | 0 | 2 | 4 |
| Calcium channel blockers (verapamil) | 0 | 1 | 0 |
| Antiarrhythmic drugs | 0 | 0 | 3 |
| Glycosides | 0 | 2 | 1 |
| Statin | 1 | 5 | 5 |
| Allopurinol | 0 | 1 | 3 |
| Diuretics | 0 | 5 | 6 |
| Aldosterone inhibitor | 0 | 1 | 1 |
| Oral anticoagulants | 0 | 7 | 9 |
| Antidepressant | 0 | 1 | 1 |
| Oral antidiabetic drugs | 0 | 3 | 2 |
| α-AR blocker | 0 | 1 | 0 |
| PPI | 1 | 7 | 9 |
| NSAID | 0 | 2 | 1 |
| Insulin | 0 | 3 | 1 |
| **Average osteopontin (ng/ml)** | 9,65 | 16,78 | 25,99 |

**Supplemental Table 2:** Patient data for cell culture, western blot and methylation-specific PCR

|  | SR (n = 27) | AF (n = 20) |
| --- | --- | --- |
| **Average age [mean, years]** | 68,8 | 72,4 |
| **Gender** |  |  |
| (Male) | 21 | 12 |
| (Female) | 6 | 8 |
| **Average BMI [kg/m<sup>2</sup>]** | 29,9 | 28,4 |
| **Disease** |  |  |
| Hypertension | 24 | 15 |
| Diabetes mellitus | 9 | 7 |
| Hyperlipidemia | 18 | 6 |
| Chronic kidney disease (GFR 40 - 90) | 7 | 7 |
| Chronic lung disease | 5 | 2 |
| Current smoking | 11 | 4 |
| Alcohol addiction | 1 | 0 |
| OSAS | 2 | 2 |
| **Diagnosis** |  |  |
| ACB | 18 | 7 |
| Valvular replacement | 11 | 16 |
| Ablation | 0 | 13 |
| **Echocardiography** |  |  |
| Ejection fraction [%] | 54,0 | 50,6 |
| LV hypertrophy | 12 | 7 |
| **Drugs** |  |  |
| ACE inhibitors | 12 | 7 |
| AT1 receptor blockers | 6 | 4 |
| β-AR blockers | 15 | 17 |
| Calcium channel blockers | 4 | 2 |
| Antiarrhythmic drugs | 0 | 2 |
| Glycosides | 0 | 9 |
| Statin | 21 | 6 |
| Allopurinol | 2 | 5 |
| Diuretics | 12 | 11 |
| Aldosterone inhibitor | 1 | 3 |
| Oral anticoagulants | 3 | 14 |
| Antidepressant | 2 | 1 |
| Oral antidiabetic drugs | 4 | 2 |
| α-AR blocker | 2 | 2 |
| PPI | 2 | 3 |
| ASS | 14 | 1 |
| Insulin | 2 | 1 |

**Supplemental Table 3:** Patient data for real-time PCR validation of the Affymetrix^®^ microarray

|  | SR (n = 7) | AF (n = 5) |
| --- | --- | --- |
| **Average age [mean, years]** | 68,0 | 69,2 |
| **Gender** |  |  |
| (Male) | 7 | 4 |
| (Female) | 0 | 1 |
| **Average BMI [kg/m<sup>2</sup>]** | 28,0 | 30,2 |
| **Disease** |  |  |
| Hypertension | 6 | 5 |
| Diabetes mellitus | 1 | 1 |
| Hyperlipidemia | 6 | 4 |
| Chronic kidney disease (GFR 40 - 90) | 2 | 2 |
| Chronic lung disease | 0 | 1 |
| Current smoking | 3 | 3 |
| Epilepsy | 1 | 0 |
| OSAS | 1 | 0 |
| **Diagnosis** |  |  |
| ACB | 7 | 3 |
| Valvular replacement | 2 | 5 |
| Ablation | 0 | 3 |
| **Echocardiography** |  |  |
| Ejection fraction [%] | 47,3 | 36,3 |
| LV hypertrophy | 2 | 2 |
| **Drugs** |  |  |
| ACE inhibitors | 5 | 4 |
| AT1 receptor blockers | 2 | 1 |
| β-AR blockers | 6 | 5 |
| Nitrates | 1 | 1 |
| Calcium channel blockers | 2 | 3 |
| Antiarrhythmic drugs | 0 | 1 |
| Glycosides | 1 | 2 |
| Statin | 6 | 4 |
| Diuretics | 2 | 4 |
| Aldosterone inhibitor | 0 | 1 |
| Oral anticoagulants | 1 | 3 |
| Antidepressant | 1 | 1 |
| Oral antidiabetic drugs | 1 | 1 |
| PPI | 2 | 1 |
| ASS | 4 | 3 |

**Supplemental table 4:**
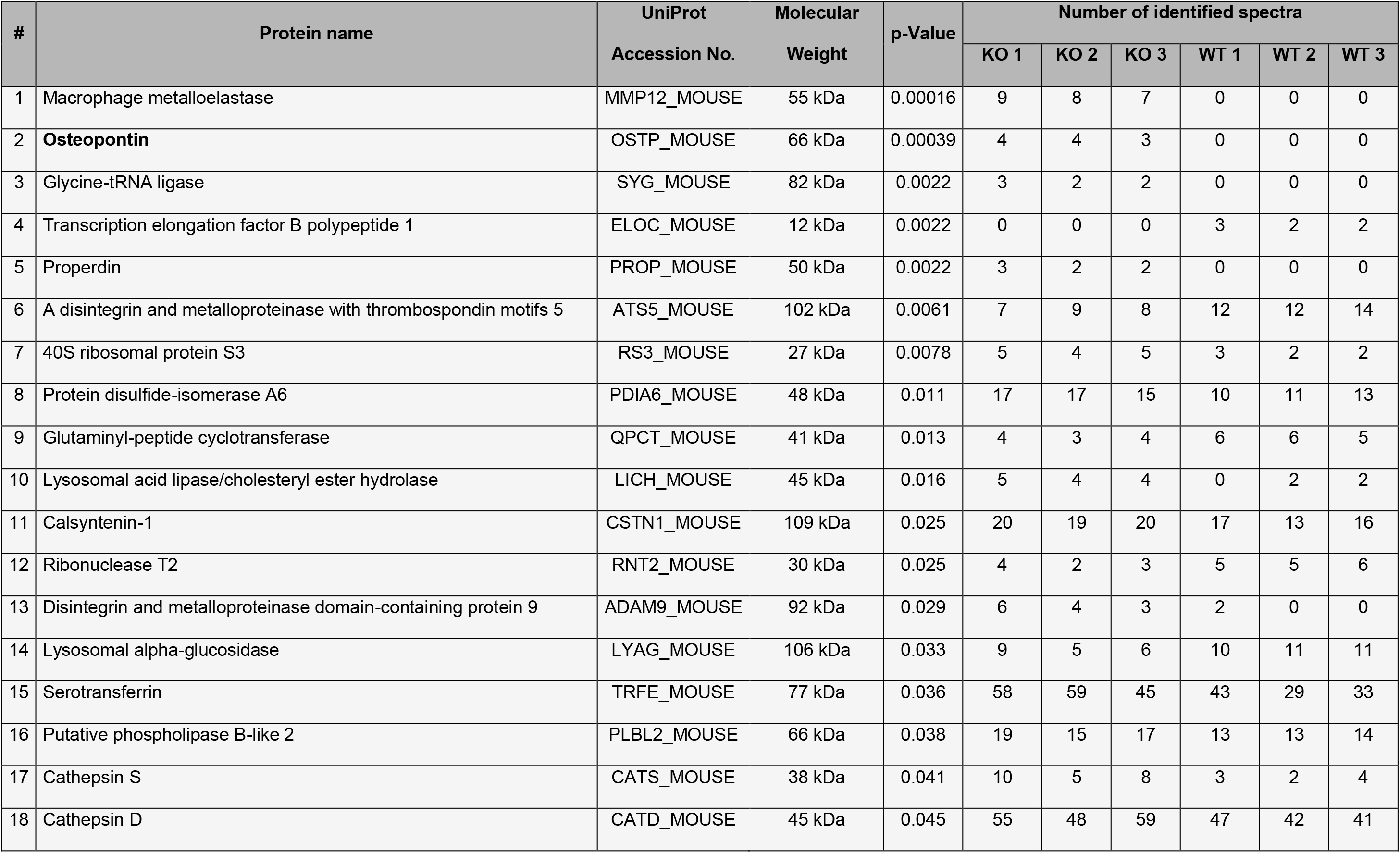
Significantly differentially regulated proteins from the fibroblast secretome analysis

## References

Abonnenc M, Nabeebaccus AA, Mayr U, Barallobre-Barreiro J, Dong X, Cuello F, Sur S, Drozdov I, Langley SR, Lu R, Stathopoulou K, Didangelos A, Yin X, Zimmermann W-H, Shah AM, Zampetaki A & Mayr M (2013) Extracellular matrix secretion by cardiac fibroblasts: role of microRNA-29b and microRNA-30c. Circ. Res. 113: 1138–1147

Ayrapetov MK, Xu C, Sun Y, Zhu K, Parmar K, D’Andrea AD & Price BD (2011) Activation of Hif1α by the prolylhydroxylase inhibitor dimethyoxalyglycine decreases radiosensitivity. PLoS ONE 6: e26064

Baum J & Duffy HS (2011) Fibroblasts and myofibroblasts: what are we talking about? J. Cardiovasc. Pharmacol. 57: 376–379

Bechtel W, McGoohan S, Zeisberg EM, Müller GA, Kalbacher H, Salant DJ, Müller CA, Kalluri R & Zeisberg M (2010) Methylation determines fibroblast activation and fibrogenesis in the kidney. Nat. Med. 16: 544–550

Benetatos L, Dasoula A, Hatzimichael E, Syed N, Voukelatou M, Dranitsaris G, Bourantas KL & Crook T (2011) Polo-like kinase 2 (SNK/PLK2) is a novel epigenetically regulated gene in acute myeloid leukemia and myelodysplastic syndromes: genetic and epigenetic interactions. Ann. Hematol. 90: 1037–1045

Bolger AM, Lohse M & Usadel B (2014) Trimmomatic: a flexible trimmer for Illumina sequence data. Bioinformatics 30: 2114–2120

Boos CJ, Anderson RA & Lip GYH (2006) Is atrial fibrillation an inflammatory disorder? Eur Heart J 27: 136–149

Calvo D, Filgueiras-Rama D & Jalife J (2018) Mechanisms and Drug Development in Atrial Fibrillation. Pharmacol. Rev. 70: 505–525

Childs BG, Durik M, Baker DJ & van Deursen JM (2015) Cellular senescence in aging and age-related disease: from mechanisms to therapy. Nat. Med. 21: 1424–1435

Chung MK, Martin DO, Sprecher D, Wazni O, Kanderian A, Carnes CA, Bauer JA, Tchou PJ, Niebauer MJ, Natale A & Van Wagoner DR (2001) C-reactive protein elevation in patients with atrial arrhythmias: inflammatory mechanisms and persistence of atrial fibrillation. Circulation 104: 2886–2891

Collado M, Blasco MA & Serrano M (2007) Cellular senescence in cancer and aging. Cell 130: 223–233

Coppé J-P, Patil CK, Rodier F, Sun Y, Muñoz DP, Goldstein J, Nelson PS, Desprez P-Y & Campisi J (2008) Senescence-associated secretory phenotypes reveal cell-nonautonomous functions of oncogenic RAS and the p53 tumor suppressor. PLoS Biol. 6: 2853–2868

Deng S, Wang H, Jia C, Zhu S, Chu X, Ma Q, Wei J, Chen E, Zhu W, Macon CJ, Jayaweera DT, Dykxhoorn DM & Dong C (2017) MicroRNA-146a Induces Lineage-Negative Bone Marrow Cell Apoptosis and Senescence by Targeting Polo-Like Kinase 2 Expression. Arterioscler. Thromb. Vasc. Biol. 37: 280–290

Dobin A, Davis CA, Schlesinger F, Drenkow J, Zaleski C, Jha S, Batut P, Chaisson M & Gingeras TR (2013) STAR: ultrafast universal RNA-seq aligner. Bioinformatics 29: 15–21

El-Armouche A, Wittköpper K, Degenhardt F, Weinberger F, Didié M, Melnychenko I, Grimm M, Peeck M, Zimmermann WH, Unsöld B, Hasenfuss G, Dobrev D & Eschenhagen T (2008) Phosphatase inhibitor-1-deficient mice are protected from catecholamine-induced arrhythmias and myocardial hypertrophy. Cardiovasc. Res. 80: 396–406

El-Tanani MK, Campbell FC, Kurisetty V, Jin D, McCann M & Rudland PS (2006) The regulation and role of osteopontin in malignant transformation and cancer. Cytokine Growth Factor Rev. 17: 463–474

Francia P, Adduci C, Semprini L, Borro M, Ricotta A, Sensini I, Santini D, Caprinozzi M, Balla C, Simmaco M & Volpe M (2014) Osteopontin and galectin-3 predict the risk of ventricular tachycardia and fibrillation in heart failure patients with implantable defibrillators. J. Cardiovasc. Electrophysiol. 25: 609–616

Geschka S, Kretschmer A, Sharkovska Y, Evgenov OV, Lawrenz B, Hucke A, Hocher B & Stasch J-P (2011) Soluble guanylate cyclase stimulation prevents fibrotic tissue remodeling and improves survival in salt-sensitive Dahl rats. PLoS ONE 6: e21853

Gramley F, Lorenzen J, Jedamzik B, Gatter K, Koellensperger E, Munzel T & Pezzella F (2010) Atrial fibrillation is associated with cardiac hypoxia. Cardiovasc. Pathol. 19: 102–111

Gu J, Wang S, Guo H, Tan Y, Liang Y, Feng A, Liu Q, Damodaran C, Zhang Z, Keller BB, Zhang C & Cai L (2018) Inhibition of p53 prevents diabetic cardiomyopathy by preventing early-stage apoptosis and cell senescence, reduced glycolysis, and impaired angiogenesis. Cell Death Dis 9: 82

Gu Z, Eils R & Schlesner M (2016) Complex heatmaps reveal patterns and correlations in multidimensional genomic data. Bioinformatics 32: 2847–2849

Güneş HM, Babur Güler G, Güler E, Demir GG, Kızılırmak Yılmaz F, Omaygenç MO, İstanbullu Tosun A, Akgün T, Boztosun B & Kılıçarslan F (2017) Relationship between serum osteopontin level and atrial fibrillation recurrence in patients undergoing cryoballoon catheter ablation. Turk Kardiyol Dern Ars 45: 26–32

Hadi HA, Alsheikh-Ali AA, Mahmeed WA & Suwaidi JMA (2010) Inflammatory cytokines and atrial fibrillation: current and prospective views. J Inflamm Res 3: 75–97

Harada M, Van Wagoner DR & Nattel S (2015) Role of inflammation in atrial fibrillation pathophysiology and management. Circ. J. 79: 495–502

Heijman J, Guichard J-B, Dobrev D & Nattel S (2018) Translational Challenges in Atrial Fibrillation. Circ. Res. 122: 752–773

Hickey FB, England K & Cotter TG (2005) Bcr-Abl regulates osteopontin transcription via Ras, PI-3K, aPKC, Raf-1, and MEK. J Leukoc Biol 78: 289–300

İçer MA & Gezmen-Karadağ M (2018) The multiple functions and mechanisms of osteopontin. Clin. Biochem.

Inglis KJ, Chereau D, Brigham EF, Chiou S-S, Schöbel S, Frigon NL, Yu M, Caccavello RJ, Nelson S, Motter R, Wright S, Chian D, Santiago P, Soriano F, Ramos C, Powell K, Goldstein JM, Babcock M, Yednock T, Bard F, et al (2009) Polo-like kinase 2 (PLK2) phosphorylates alpha-synuclein at serine 129 in central nervous system. J. Biol. Chem. 284: 2598–2602

Krtolica A, Parrinello S, Lockett S, Desprez PY & Campisi J (2001) Senescent fibroblasts promote epithelial cell growth and tumorigenesis: a link between cancer and aging. Proc. Natl. Acad. Sci. U.S.A. 98: 12072–12077

Lee KJ, Hoe H-S & Pak DT (2011a) Plk2 Raps up Ras to subdue synapses. Small GTPases 2: 162–166

Lee KJ, Lee Y, Rozeboom A, Lee J-Y, Udagawa N, Hoe H-S & Pak DTS (2011b) Requirement for Plk2 in orchestrated ras and rap signaling, homeostatic structural plasticity, and memory. Neuron 69: 957–973

Li J, Ma W, Wang P, Hurley PJ, Bunz F & Hwang PM (2014) Polo-like kinase 2 activates an antioxidant pathway to promote the survival of cells with mitochondrial dysfunction. Free Radic. Biol. Med. 73: 270–277

Liao Y, Smyth GK & Shi W (2013) The Subread aligner: fast, accurate and scalable read mapping by seed-and-vote. Nucleic Acids Res. 41: e108

López B, González A, Lindner D, Westermann D, Ravassa S, Beaumont J, Gallego I, Zudaire A, Brugnolaro C, Querejeta R, Larman M, Tschöpe C & Díez J (2013) Osteopontin-mediated myocardial fibrosis in heart failure: a role for lysyl oxidase? Cardiovasc. Res. 99: 111–120

Love MI, Huber W & Anders S (2014) Moderated estimation of fold change and dispersion for RNA-seq data with DESeq2. Genome Biol. 15: 550

Ma S, Charron J & Erikson RL (2003) Role of Plk2 (Snk) in mouse development and cell proliferation. Mol. Cell. Biol. 23: 6936–6943

Mochizuki M, Lorenz V, Ivanek R, Della Verde G, Gaudiello E, Marsano A, Pfister O & Kuster GM (2017) Polo-Like Kinase 2 is Dynamically Regulated to Coordinate Proliferation and Early Lineage Specification Downstream of Yes-Associated Protein 1 in Cardiac Progenitor Cells. J Am Heart Assoc 6:

Nattel S, Burstein B & Dobrev D (2008) Atrial remodeling and atrial fibrillation: mechanisms and implications. Circ Arrhythm Electrophysiol 1: 62–73

Parrinello S, Coppe J-P, Krtolica A & Campisi J (2005) Stromal-epithelial interactions in aging and cancer: senescent fibroblasts alter epithelial cell differentiation. J. Cell. Sci. 118: 485–496

Plumb JA, Strathdee G, Sludden J, Kaye SB & Brown R (2000) Reversal of drug resistance in human tumor xenografts by 2’-deoxy-5-azacytidine-induced demethylation of the hMLH1 gene promoter. Cancer Res. 60: 6039–6044

Poulet C, Künzel Stephan, Büttner E, Lindner D, Westermann D & Ravens U (2016) Altered physiological functions and ion currents in atrial fibroblasts from patients with chronic atrial fibrillation. Physiol Rep 4:

R Core Team (2014) R: A language and environment for statistical computing. R Foundation for Statistical Computing, Vienna, Austria. URL http://www.R-project.org/.

Robinson CM, Neary R, Levendale A, Watson CJ & Baugh JA (2012) Hypoxia-induced DNA hypermethylation in human pulmonary fibroblasts is associated with Thy-1 promoter methylation and the development of a pro-fibrotic phenotype. Respir. Res. 13: 74

Rubiś P, Wiśniowska-Śmiał k S, Dziewięcka E, Rudnicka-Sosin L, Kozanecki A & Podolec P (2018) Prognostic value of fibrosis-related markers in dilated cardiomyopathy: A link between osteopontin and cardiovascular events. Adv Med Sci 63: 160–166

Rudolph V, Andrié RP, Rudolph TK, Friedrichs K, Klinke A, Hirsch-Hoffmann B, Schwoerer AP, Lau D, Fu X, Klingel K, Sydow K, Didié M, Seniuk A, von Leitner E-C, Szoecs K, Schrickel JW, Treede H, Wenzel U, Lewalter T, Nickenig G, et al (2010) Myeloperoxidase acts as a profibrotic mediator of atrial fibrillation. Nat. Med. 16: 470–474

Sandner P, Berger P & Zenzmaier C (2017) The Potential of sGC Modulators for the Treatment of Age-Related Fibrosis: A Mini-Review. Gerontology 63: 216–227

Schott S, Wimberger P, Klink B, Grützmann K, Puppe J, Wauer US, Klotz DM, Schröck E & Kuhlmann JD (2017) The conjugated antimetabolite 5-FdU-ECyd and its cellular and molecular effects on platinum-sensitive vs. -resistant ovarian cancer cells in vitro. Oncotarget 8: 76935–76948

Syed N, Smith P, Sullivan A, Spender LC, Dyer M, Karran L, O’Nions J, Allday M, Hoffmann I, Crawford D, Griffin B, Farrell PJ & Crook T (2006) Transcriptional silencing of Polo-like kinase 2 (SNK/PLK2) is a frequent event in B-cell malignancies. Blood 107: 250–256

Tallquist MD & Molkentin JD (2017) Redefining the identity of cardiac fibroblasts. Nat Rev Cardiol 14: 484–491

Thomas SM, DeMarco M, D’Arcangelo G, Halegoua S & Brugge JS (1992) Ras is essential for nerve growth factor-and phorbol ester-induced tyrosine phosphorylation of MAP kinases. Cell 68: 1031–1040

Verma A, Wazni OM, Marrouche NF, Martin DO, Kilicaslan F, Minor S, Schweikert RA, Saliba W, Cummings J, Burkhardt JD, Bhargava M, Belden WA, Abdul-Karim A & Natale A (2005) Pre-existent left atrial scarring in patients undergoing pulmonary vein antrum isolation: an independent predictor of procedural failure. J. Am. Coll. Cardiol. 45: 285–292

Vílchez JA, Roldán V, Hernández-Romero D, Valdés M, Lip GYH & Marín F (2014) Biomarkers in atrial fibrillation: an overview. Int. J. Clin. Pract. 68: 434–443

Watanabe T, Takeishi Y, Hirono O, Itoh M, Matsui M, Nakamura K, Tamada Y & Kubota I (2005) C-reactive protein elevation predicts the occurrence of atrial structural remodeling in patients with paroxysmal atrial fibrillation. Heart Vessels 20: 45–49

Watson CJ, Collier P, Tea I, Neary R, Watson JA, Robinson C, Phelan D, Ledwidge MT, McDonald KM, McCann A, Sharaf O & Baugh JA (2014) Hypoxia-induced epigenetic modifications are associated with cardiac tissue fibrosis and the development of a myofibroblast-like phenotype. Hum Mol Genet 23: 2176–2188

Xie Z, Singh M & Singh K (2004) ERK1/2 and JNKs, but not p38 kinase, are involved in reactive oxygen species-mediated induction of osteopontin gene expression by angiotensin II and interleukin-1β in adult rat cardiac fibroblasts. J. Cell. Physiol. 198: 399–407

Yin X, Bern M, Xing Q, Ho J, Viner R & Mayr M (2013) Glycoproteomic analysis of the secretome of human endothelial cells. Mol. Cell Proteomics 12: 956–978

Zhao H, Wang W, Zhang J, Liang T, Fan G-P, Wang Z-W, Zhang P-D, Wang X & Zhang J (2016) Inhibition of osteopontin reduce the cardiac myofibrosis in dilated cardiomyopathy via focal adhesion kinase mediated signaling pathway. Am J Transl Res 8: 3645–3655

Zhu F, Li Y, Zhang J, Piao C, Liu T, Li H-H & Du J (2013) Senescent cardiac fibroblast is critical for cardiac fibrosis after myocardial infarction. PLoS ONE 8: e74535

